# Acentrosomal spindle assembly and stability in *C. elegans* oocytes requires a kinesin-12 non-motor microtubule interaction domain

**DOI:** 10.1101/2021.06.17.448874

**Authors:** Ian D. Wolff, Jeremy A. Hollis, Sarah M. Wignall

## Abstract

During the meiotic divisions in oocytes, microtubules are sorted and organized by motor proteins to generate a bipolar spindle in the absence of centrosomes [1]. In most organisms, kinesin-5 family members crosslink and slide microtubules to generate outward force that promotes acentrosomal spindle bipolarity [2–7]. However, the mechanistic basis for how other kinesin families act on acentrosomal spindles has not been explored. We investigated this question in *C. elegans* oocytes, where kinesin-5 is not required to generate outward force [8]. Instead, the kinesin-12 family motor KLP-18 performs this function [9–12]. KLP-18 acts with adaptor protein MESP-1 (<u>me</u>iotic <u>sp</u>indle <u>1</u>) to sort microtubule minus ends to the periphery of a microtubule array, where they coalesce into spindle poles [12]. If either of these proteins is depleted, outward sorting of microtubules is lost and minus ends converge to form a monoaster. Here we use a combination of *in vitro* biochemical assays and *in vivo* mutant analysis to provide insight into the mechanism by which these proteins collaborate to promote acentrosomal spindle assembly. We identify a microtubule binding site on the C-terminal stalk of KLP-18 and demonstrate that a direct interaction between the KLP-18 stalk and MESP-1 activates non-motor microtubule binding. We also provide evidence that this C-terminal domain is required for KLP-18 activity during spindle assembly and show that KLP-18 is continuously required to maintain spindle bipolarity. This study thus provides new insight into the construction and maintenance of the oocyte acentrosomal spindle as well as into kinesin-12 mechanism and regulation.

## RESULTS

### The KLP-18 coiled-coil stalk domain contains a regulated microtubule binding site

In mammalian mitosis, kinesin-12 (Kif15) has been shown to be able to promote spindle bipolarity when kinesin-5 is inhibited, suggesting that these motors both provide outward forces that serve to separate spindle poles [13–15]. Kinesin-12 family motors contain a globular N-terminal motor domain that can walk along microtubules in a plus end-directed fashion. However, in order to crosslink and sort microtubules within the spindle, these motors must bind a second microtubule. Multiple microtubule cross-linking strategies have been proposed for Kif15: 1) Kif15 forms a homotetramer with antipolar motor domains, allowing both ends of the complex to bind to microtubules [16–19], and 2) a non-motor microtubule-binding site in the middle of the stalk domain mediates binding to a second microtubule [20, 21]. Since the kinesin-12 family motor KLP-18 is the major microtubule sorting kinesin in the *C. elegans* oocyte spindle, we reasoned that this would be an ideal system to investigate the mechanisms by which kinesin-12 motors promote spindle bipolarity, starting with the question of whether this motor contains multiple microtubule binding sites.

Like other kinesin-12 motors, KLP-18 contains a C-terminal stalk domain [9], so we first examined if this domain could mediate crosslinking by directly binding to microtubules. Structural prediction of the KLP-18 stalk revealed that it contains discrete coiled-coil domains (Fig 1A), similar to Kif15 [9, 20]. To test if the stalk contains a microtubule binding site, we expressed and purified the N- and C-terminal halves of the stalk (“N-stalk” and “C-stalk”; Fig 1A, S1A) and performed a microtubule co-sedimentation assay [22] (Fig 1B). After pelleting microtubules, N-stalk remained in the supernatant, while a sizable fraction of C-stalk pelleted, reflecting its ability to bind microtubules *in vitro* (Fig 1B, left). To investigate the nature of this interaction, we treated microtubules with subtilisin to cleave E-hooks, which are negatively-charged regions at the tubulin C-terminus that can promote microtubule association of proteins through an electrostatic interaction [23, 24]. We found that C-stalk bound to microtubules lacking E-hooks, but with decreased affinity (Fig 1B, right). These results are consistent with previous studies of the non-motor microtubule binding sites in Kif15 and kinesin-1 [21, 25], and suggest that an electrostatic interaction increases the affinity of KLP-18 stalk to microtubules. To further assess the microtubule binding activity of C-stalk and N-stalk, we incubated these proteins with fluorescently-labeled microtubules *in vitro* and visualized microtubule organization; during purification, both N-stalk and C-stalk elute from size exclusion chromatography as oligomeric complexes, potentially providing multiple binding sites that could facilitate microtubule bundling. Addition of C-stalk caused dramatic microtubule bundling compared to buffer alone or N-stalk (Fig 1C), confirming that C-stalk is able to bind microtubules. Together, these data show that the stalk domain of KLP-18 contains a C-terminal microtubule binding site.

**Figure 1.**
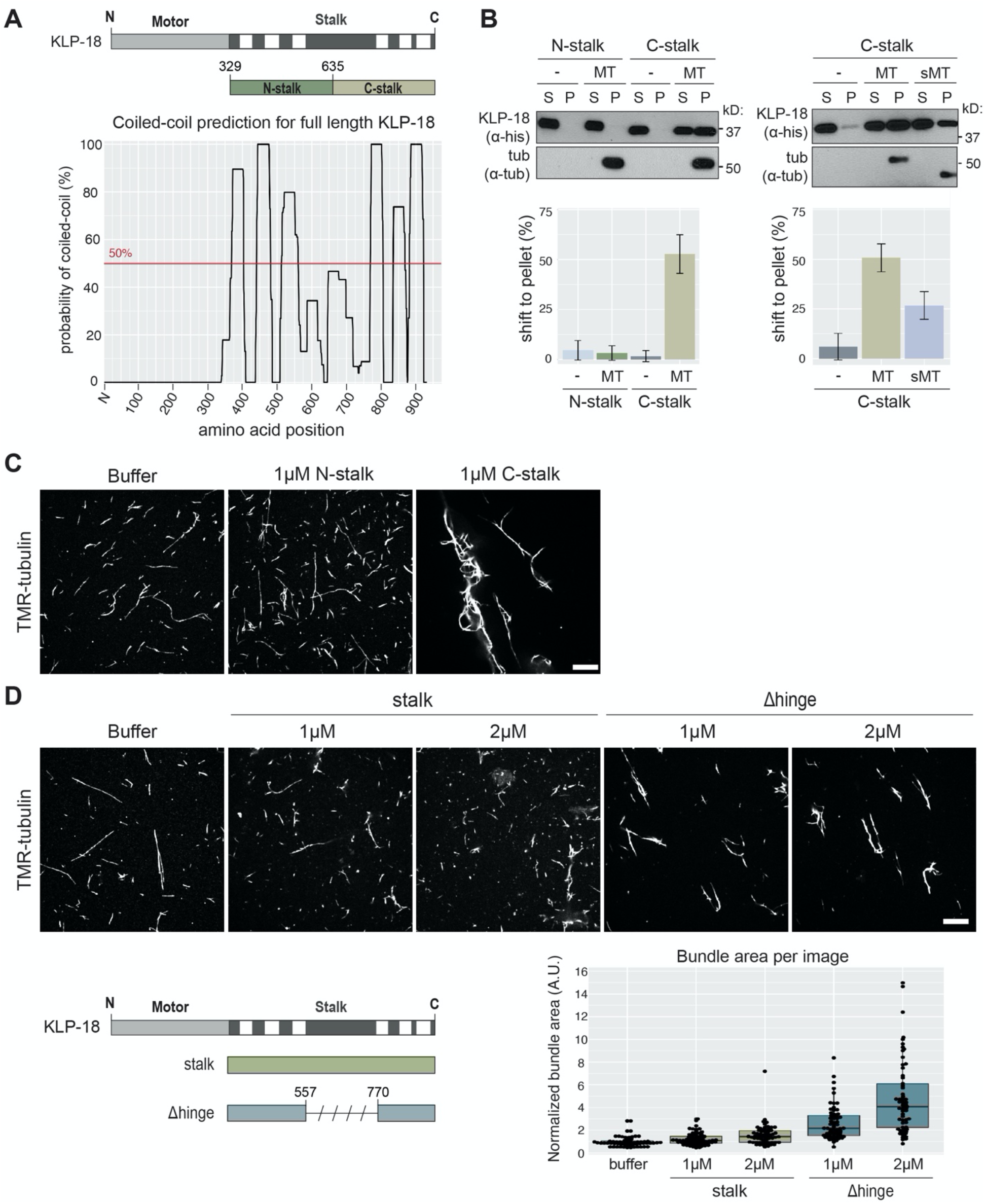
The kinesin-12/KLP-18 coiled-coil stalk domain contains a regulated microtubule binding site. a) Coiled-coil prediction software paircoil2 [62] shows discrete coiled-coil domains in the KLP-18 stalk (bottom) marked on a KLP-18 domain diagram (top, coiled-coil domains denoted in white). N- and C-stalk constructs shown relative to full stalk. b) N-stalk and C-stalk microtubule binding activity tested by microtubule co-sedimentation assay with no microtubules added (-), undigested (MT), and subtilisin-digested (sMT) microtubules. Blots show supernatant (S) and pellet (P) samples, quantification is of average shift +/- sd. N = 3 experiments for each set. c, d) Microtubule binding activity assessed as microtubule bundling ability. Representative images of TMR-microtubules incubated with buffer alone, N-stalk, and C-stalk (c) or with buffer alone, full-length stalk, and Δhinge (d). Quantification of bundling below. Boxplot represents first quartile to third quartile and the median is indicated by a horizontal line. Quantified images were acquired over 2 independent experiments. Scale bars = 10μm.

Next, we purified the full KLP-18 stalk (termed “stalk”; Fig S1A) and paradoxically found that it showed very little microtubule bundling activity (Fig 1D), suggesting that it cannot bind microtubules. Since an autoinhibitory mechanism has been proposed for mammalian Kif15, where the motor folds its stalk domain to block the non-motor microtubule binding site [20], we tested whether KLP-18 could employ a similar regulatory mechanism. The coiled-coil prediction for the KLP-18 stalk contains a region of low probability in the center of the stalk domain (Fig 1A), and we hypothesized that this region may be a flexible hinge that could fold, thus preventing the C-terminal region from binding microtubules. To test this mechanism, we purified a version of the stalk with this putative hinge deleted (termed “Δhinge”), which is predicted to be completely coiled-coil (Fig S1A, S2A, B). Consistent with an auto-inhibitory mechanism, deletion of the hinge region restored microtubule bundling activity, reflecting an ability to bind microtubules (Fig 1D). In addition, we assessed the flexibility of the KLP-18 stalk by running an MBP-stalk fusion protein (Fig S2A) through a size exclusion column in high salt (300mM) and low salt (20mM) buffers. If the KLP-18 stalk was able to fold via an electrostatic interaction, high salt would disrupt folding and lead to an extended conformation; this conformational change would be apparent in the elution volume. Indeed, in high salt, a population of MBP-stalk eluted at a lower elution volume, indicating that this sub-population contains extended molecules (Fig S2C). These results are consistent with the possibility that the KLP-18 stalk can exist in either an inactive folded state, or an unfolded state that is capable of binding to microtubules.

### The KLP-18 stalk microtubule interaction domain is essential for spindle assembly

During acentrosomal spindle assembly in *C. elegans* oocytes, KLP-18 sorts microtubule minus ends to the outside of a microtubule array, enabling the formation of multiple poles that then coalesce to form a bipolar spindle [9–12]. Given our identification of the C-terminal microtubule binding site, we hypothesized that KLP-18 might sort microtubules by binding a microtubule with its stalk and walking on a second microtubule with its motor domain. To test this hypothesis, we analyzed a set of *klp-18* mutants *in vivo*. We previously demonstrated that a mutant lacking most of the N-terminal motor domain, *klp-18(ok2519)*, formed monopolar oocyte spindles, confirming that this domain is required for KLP-18 function [12]. To assess requirements for other domains, we made use of the previously described *klp-18(or447)* temperature sensitive mutant, which contains two substitutions (V854M and G876S) in the C-terminal microtubule binding domain (Fig 2A, mutant hereafter referred to as ‘*klp-18ts’*) [11]. At the restrictive temperature of 26°C, *klp-18ts* has high embryonic lethality, defects in polar body extrusion, and aberrant chromosome dynamics, suggesting that bipolar spindles do not form [11], but this has not been directly demonstrated. In addition, we generated two new *klp-18* mutants using CRISPR: *klp-18ΔC-stalk*, a deletion of the C-terminal microtubule binding site, and *klp-18Δhinge*, a deletion of the putative hinge (Fig 2A), and we assessed spindle assembly in all three mutants.

**Figure 2:**
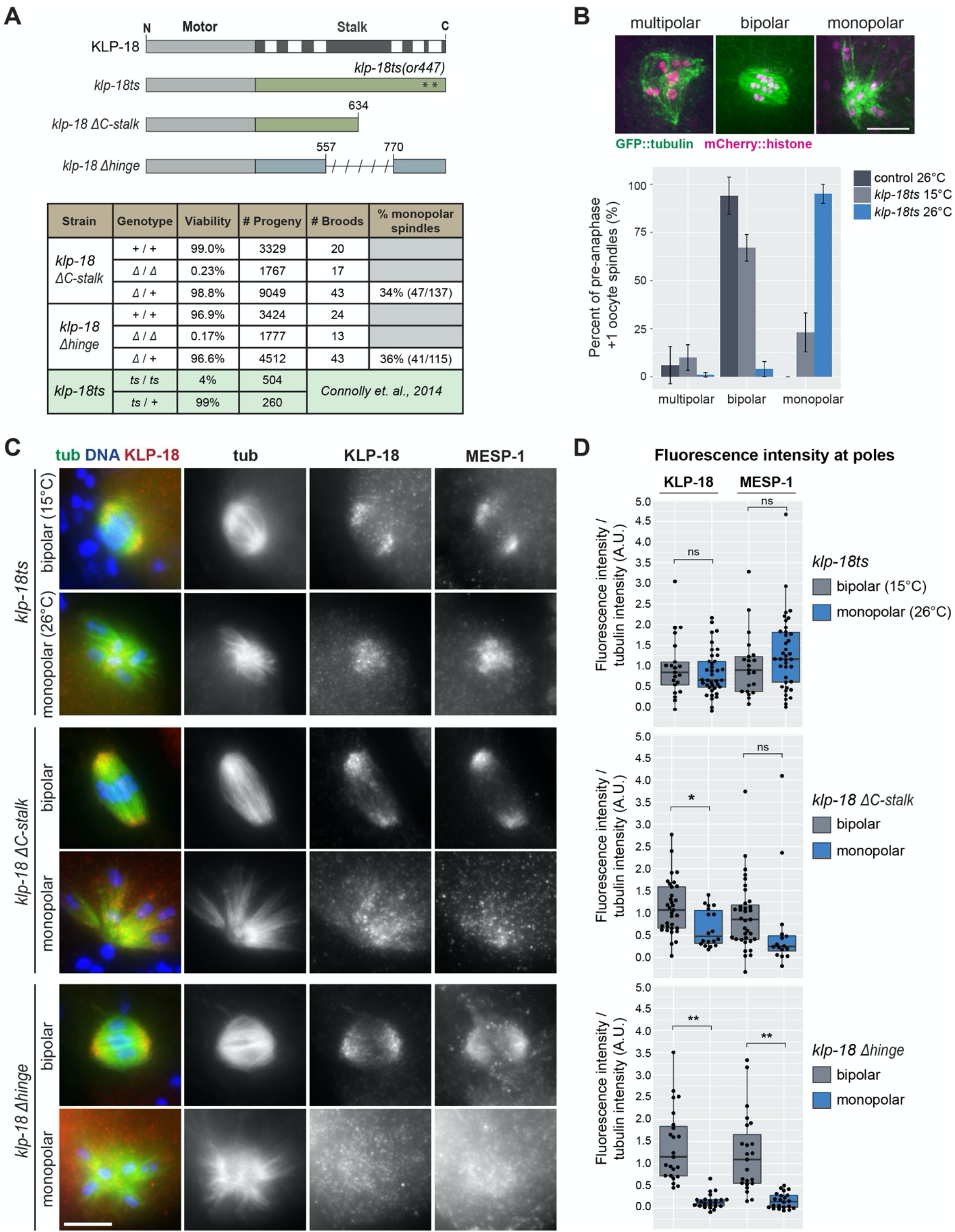
KLP-18 hinge and stalk microtubule interaction domain are essential for spindle assembly. a) Domain diagram of *klp-18* mutants used: *klp-18ts, klp-18ΔC-stalk*, and *klp-18Δhinge*. Asterisks indicate approximate location of *or447* mutations in the *klp-18ts* mutant. Characterization of progeny from heterozygous (*Δ* /+) or homozygous (*Δ* /*Δ* or +/+) *klp-18ΔC-stalk* and *klp-18Δhinge* parents. Data for *klp-18ts* was adapted from Connolly et. al., 2014 [11] and is shaded in green. b) Morphology of pre-anaphase oocyte spindles in the most recently fertilized (+1) embryo of control (dark gray) and *klp-18ts* worms expressing GFP::tubulin and mCherry::histone. During normal spindle assembly, minus ends are sorted away from chromosomes and form multiple poles (these forming spindles are denoted “multipolar”) that coalesce to form a bipolar spindle (“bipolar”) [12]. If minus ends are not sorted outwards, they converge to form a monopolar spindle (“monopolar”) [9–12]. Spindle morphology in *klp-18ts* was quantified at permissive (15°C, light gray) and restrictive (26°C, blue) temperatures. Note that spindles that had entered anaphase were not included in this graph; quantification of all +1 oocytes is shown in Figure S3A. Bars represent mean percentage +/- sd. For *klp-18ts* conditions, n = 4 experiments; for control, n = 3 experiments. Scale bar = 5μm. c) DNA (blue), tubulin (green), KLP-18 (red), and MESP-1 (not shown in merge) localization in mutant strains described in (a). Representative images of bipolar and mutant monopolar spindles are shown for each strain. Scale bar = 5μm. d) Quantification of KLP-18 and MESP-1 fluorescence intensity on the poles of bipolar (gray) and monopolar (blue) spindles in each mutant. KLP-18/MESP-1 intensity was normalized to tubulin intensity. Box represents first quartile to third quartile and the median is indicated by a horizontal line. Welch Two-Sample t-test, ns = p > 0.05, * = p < 0.05, ** = p < 0.0005.

Eggs laid by homozygous parents of both the *klp-18ΔC-stalk* and *klp-18Δhinge* mutants are largely inviable (Fig 2A, 0.23% and 0.17% viability, respectively), so we were unable to propagate homozygous mutant strains. Instead, we analyzed progeny of heterozygous worms; 25% of their progeny should be homozygous mutants. Homozygous mutants reach adulthood since they have a maternal supply of wild-type KLP-18 that enables proper development, but produce oocytes containing only the mutant form of KLP-18. Notably, about 35% of pre-anaphase oocyte spindles from the adult progeny of heterozygous parents were monopolar in both strains (Fig 2A), demonstrating that deleting either the putative hinge or the C-terminal domain affects KLP-18 function. To similarly quantify spindle defects in the *klp-18ts* strain, we shifted worms to the restrictive temperature for 1 hour, thus disrupting KLP-18 before the initiation of spindle assembly. We then quantified spindle morphology in the most recently fertilized oocyte; although by this stage some control spindles had progressed to anaphase (Fig S3A), many were either forming or bipolar (Fig 2B, S3A), enabling us to assess effects on spindle formation. In *klp-18ts* oocytes at the permissive temperature (15°C), we found that 22% of pre-anaphase spindles (23/105) were monopolar. This indicates that KLP-18 function is partially compromised, consistent with previously-reported embryonic lethality at this temperature [11]. Upon shift to the restrictive temperature, the percentage of pre-anaphase spindles that were monopolar increased to 94% (124/132) (Fig 2B). As expected, the poles of monopolar spindles in *klp-18ts, klp-18ΔC-stalk*, and *klp-18Δhinge* oocytes are marked by the minus-end marker ASPM-1 [10, 26], phenocopying *klp-18(RNAi)* and confirming that microtubule sorting was aberrant (Fig S3B). However, ASPM-1 also localized to some microtubule ends on the outside of the aster in 24/30 *klp-18ts* monopolar spindles analyzed (Fig S3B, arrowheads), suggesting that *klp-18ts* may allow weak sorting activity that enables some minus ends to be pushed outwards, even at the restrictive temperature. Supporting this idea, ∼5% of pre-anaphase spindles (7/132) are bipolar at 26°C (Fig 2B).

Although our results suggest that the C-terminal microtubule binding and hinge domains are required for KLP-18 function, it is also possible that these mutations merely destabilize the KLP-18 protein. However, we did not detect a decrease in KLP-18 abundance in *klp-18ts* worms at 15°C or 26°C by Western blot compared to control worms (Fig S3C), and we found that the KLP-18ΔC-stalk and KLP-18Δhinge mutant proteins are expressed (Fig S3D), suggesting that the phenotypes in the *klp-18* mutants are not due to loss of KLP-18 protein.

In both *Xenopus* and mammals, the adaptor protein TPX2 is required for targeting kinesin-12 to the spindle [16, 18, 19, 27–29], and our previous work indicates that MESP-1 performs this kinesin-12 targeting function in *C. elegans:* KLP-18 and MESP-1 colocalize on spindle microtubules, are found in a complex in worm extract, have identical depletion phenotypes, and are interdependent for localization [12]. Therefore, to better understand the phenotype of the *klp-18* mutants, we investigated whether targeting of the KLP-18/MESP-1 complex to microtubules was affected. We quantified the fluorescence intensity of each protein at spindle poles (either an average of the two bipolar poles or the one monopolar pole) and normalized this value to the average fluorescence intensity of tubulin. In *klp-18ts* oocytes there was no significant difference between KLP-18 and MESP-1 localization to the monopole at the restrictive temperature compared to localization on the bipolar poles at the permissive temperature (Fig 2C and D, top), suggesting that the KLP-18/MESP-1 complex is able to localize to spindle microtubules but is inactivated by the temperature shift, possibly through disruption of the C-terminal microtubule binding domain that contains the two *klp-18ts* mutations. In *klp-18ΔC-stalk* oocytes there was persistent but decreased localization of KLP-18 and MESP-1 to monopoles in most oocytes, indicating that these proteins are able to target to spindle poles with decreased efficiency (Fig 2C and D, middle). Finally, we saw a striking decrease of KLP-18 and MESP-1 localization in *klp-18Δhinge* oocytes, suggesting that the “hinge” region is essential for spindle localization of either of these proteins (Fig 2C and D, bottom). Altogether, these results show that the C-terminal microtubule binding site and the putative hinge region we identified *in vitro* are essential for KLP-18 activity and spindle assembly *in vivo*.

### KLP-18 is targeted to microtubules through a direct interaction with microtubule associated protein MESP-1

Next, we aimed to better understand the contribution of MESP-1 to KLP-18 function; MESP-1 is required for KLP-18 spindle localization, but how MESP-1 regulates KLP-18 is not known. Considering that MESP-1 can localize to monopolar spindle microtubules in the *klp-18ΔC-stalk* and *klp-18ts* strains, in which C-terminal microtubule binding is disrupted, but does not localize in the *klp-18Δhinge* mutant, we hypothesized that MESP-1 may bind to a region overlapping with the hinge. To test this *in vitro*, we first attempted to express GST-MESP-1, but this protein degraded significantly during purification (Fig S1B). This suggests that MESP-1 is unstable, consistent with the prediction that portions of MESP-1 are disordered (Fig S1C) similar to TPX2 [22, 30, 31]. Therefore, we switched to an MBP tag [32–34], which increased MESP-1 stability (Fig S1B). We incubated purified MBP-MESP-1 with the N- and C-terminal KLP-18 stalk truncations individually and added amylose resin to retrieve MBP-MESP-1. We found that N-stalk was present in the eluted fraction but C-stalk was not, indicating that MESP-1 binds to the N-terminal half of the KLP-18 stalk (Fig 3A). However, this interaction appears to be weak, because only a small fraction of N-stalk was pulled out by MBP-MESP-1; this is similar to what has been reported for the interaction between mammalian TPX2 and Kif15 [16]. Importantly, this N-terminal MESP-1 binding region overlaps significantly with the hinge region (Fig S4A), perhaps explaining why MESP-1 localization is decreased in *klp-18Δhinge* spindles.

**Figure 3.**
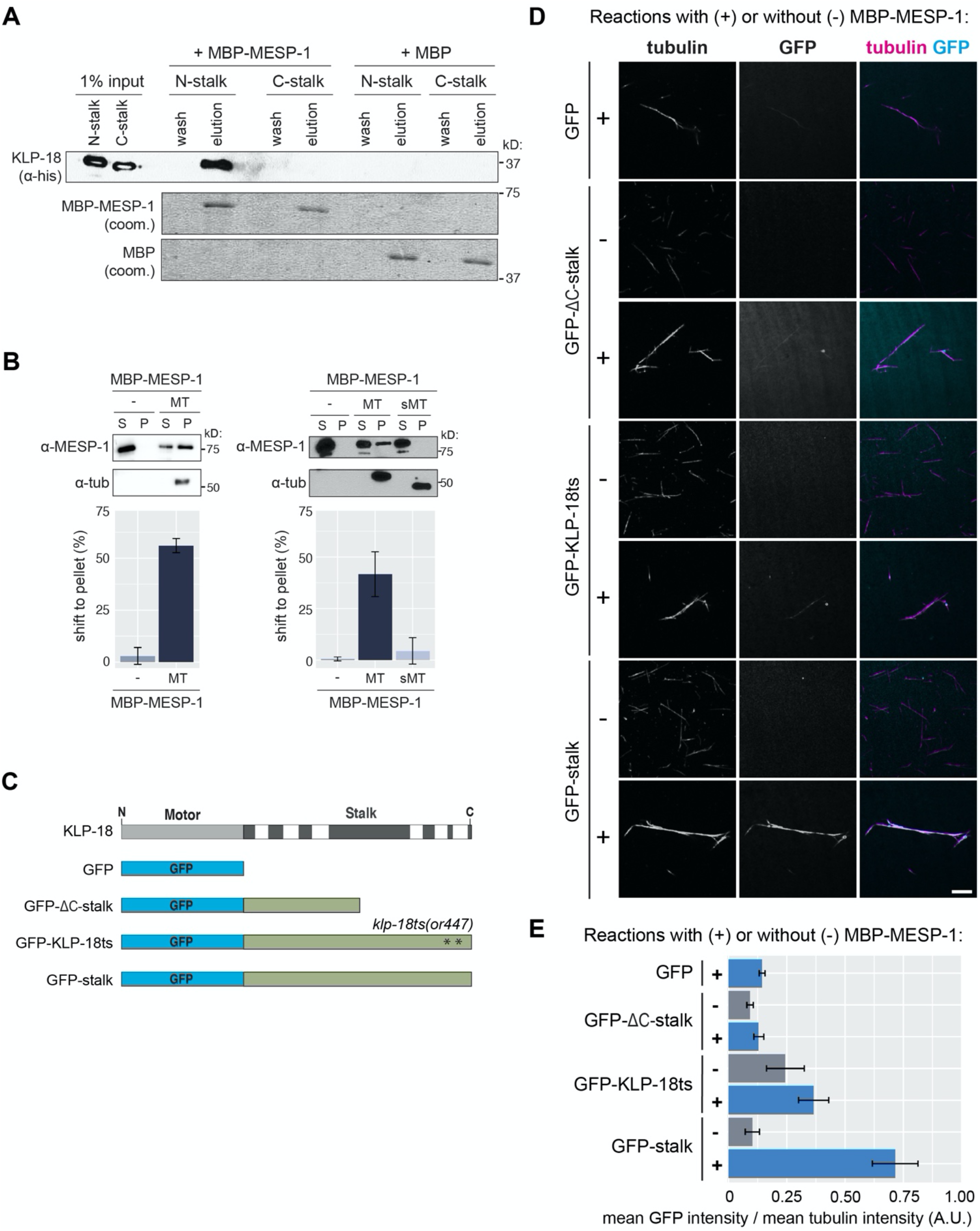
KLP-18 C-terminal domain is essential for microtubule binding *in vitro* and requires a direct interaction with microtubule associated protein MESP-1. a) MBP-MESP-1 direct interaction with N-stalk or C-stalk was tested by MBP pulldown. Reactions using MBP-MESP-1 or MBP as bait are shown. KLP-18 truncations visualized on Western blot with anti-His antibody; MBP and MBP-MESP-1 visualized by Coomassie stain. b) Microtubule co-sedimentation assay for MBP-MESP-1 with no microtubules added (-), undigested (MT), and subtilisin-digested (sMT) microtubules. Representative blots show supernatant (S) and pellet (P) samples, quantification is of average shift +/- sd. n = 2 experiments for non-subtilisin experiment, n = 3 experiments for subtilisin experiment. c) Schematic of GFP tagged proteins used in (d) and (e). d) KLP-18 recruitment to microtubules was tested by visualization of GFP tagged stalk proteins on TMR-microtubules. Representative images with (+) and without (-) MBP-MESP-1 added to the indicated GFP-tagged proteins are shown for each condition. Scale bar = 5μm. e) Quantification of mean GFP intensity overlayed on the microtubule normalized to mean tubulin intensity. Bars represent mean percentage +/- sd for each experiment over n = 3 experiments.

Next, we tested the microtubule binding ability of MESP-1 to begin to clarify the role of MESP-1 in activating KLP-18 and targeting it to microtubules. MBP-MESP-1 pelleted with microtubules in a co-sedimentation assay (Fig 3B) and was able to bundle microtubules (Fig S1D), confirming that MESP-1 is a microtubule binding protein. In addition, we found that MESP-1’s ability to bind microtubules was greatly diminished when we used subtilisin-digested microtubules (Fig 3B, right). Thus, MESP-1 is a microtubule associated protein, similar to TPX2. However, TPX2 binds along the microtubule lattice and does not require tubulin E-hooks [30, 35], suggesting that MESP-1 employs a different microtubule binding mechanism.

Considering that MESP-1 can bind to both KLP-18 and microtubules, we hypothesized that MESP-1 may recruit KLP-18 to microtubules directly. To test this model, we purified GFP-tagged versions of the full length KLP-18 stalk (termed “GFP-stalk”), KLP-18ΔC-stalk (“GFP-KLP-18ΔC-stalk”), and a version of the stalk containing the two mutations present in the *klp-18ts* strain (“GFP-KLP-18ts”), then quantified GFP localization to microtubules in the presence and absence of MESP-1 (Fig 3C-E). We attempted to purify a GFP-KLP-18Δhinge protein but it was unstable and did not yield consistent results. In these experiments, GFP-stalk did not localize to microtubules when incubated alone (Fig 3D, E), consistent with our model of KLP-18 auto-inhibition. However, we observed strong localization of GFP-stalk to microtubules in the presence of MBP-MESP-1 (Fig 3D, E), showing that MESP-1 is sufficient to enable KLP-18 microtubule binding. Next, we analyzed GFP-ΔC-stalk, which contains the N-terminal MESP-1 binding region (Fig 3A) but lacks the C-terminal microtubule binding domain (Fig 1B). We did not see any enrichment of GFP-ΔC-stalk onto microtubules with or without MBP-MESP-1, showing that MESP-1 alone is not sufficient to target KLP-18 to microtubules and that both the C-terminal microtubule binding domain and MESP-1 are required. Finally, we tested if the substitution mutations present in *klp-18ts* affect the microtubule binding ability of the stalk. We quantified a ∼2-fold decrease in microtubule-associated GFP-KLP-18ts with MBP-MESP-1 compared to wild type GFP-stalk with MBP-MESP-1, suggesting that the *in vivo* phenotype that we observe in the *klp-18ts* mutant is a result of decreased microtubule binding ability of the C-terminal domain.

### KLP-18 activity is essential to maintain spindle bipolarity

Given the importance of KLP-18 in spindle assembly, we next asked if KLP-18 is also necessary to maintain spindle bipolarity. In human oocytes, the acentrosomal meiotic divisions proceed through an extended bipolar stage that can become unstable, and this instability is correlated with errors in chromosome segregation that result in aneuploidy [36]. Therefore, it is important to understand the molecular mechanisms that maintain acentrosomal spindle stability.

Kinesin-5 is essential for acentrosomal spindle maintenance in mouse and *Drosophila* oocytes [5, 37, 38], but whether kinesin-12 can similarly maintain spindle bipolarity has not been tested in oocytes of any system. The rapid *klp-18ts* temperature sensitive mutant allowed us to address this question by inactivating KLP-18 function after spindles have already formed. To this end, we induced metaphase I arrest by depleting anaphase promoting complex (APC) component EMB-30 using RNAi [39]. Since the *C. elegans* germline is organized in a production-line fashion, oocytes continue to be fertilized despite this depletion, and each forms a spindle that arrests at Metaphase I; this leads to a buildup of bipolar spindles in the germ line (with the most-recently fertilized oocyte containing the spindle that was most recently formed). In *klp-18ts; emb-30(RNAi)* worms at the permissive temperature (15°C) and control *emb-30(RNAi)* worms at 26°C, at least 60% of the two most recently arrested oocyte spindles were bipolar (these embryos are denoted +1 and +2; Fig 4A). In contrast, when *klp-18ts; emb-30(RNAi)* worms were shifted to the restrictive temperature (26°C), the majority of oocyte spindles were monopolar (93% (+/- 6%) in +1 and 83% (+/- 12%) in +2 embryos). Moreover, we also observed monopolar spindles at positions beyond the +2 position in the germline (Fig 4D). Under the conditions of our temperature shift, spindles in the +1 position may have formed after KLP-18 was inactivated, but spindles in oocytes that had been arrested longer should have established bipolarity before KLP-18 inactivation. These results suggest that KLP-18 is not only required to sort microtubule minus ends outwards during spindle assembly, but also provides outward force that is required to maintain spindle bipolarity.

**Figure 4:**
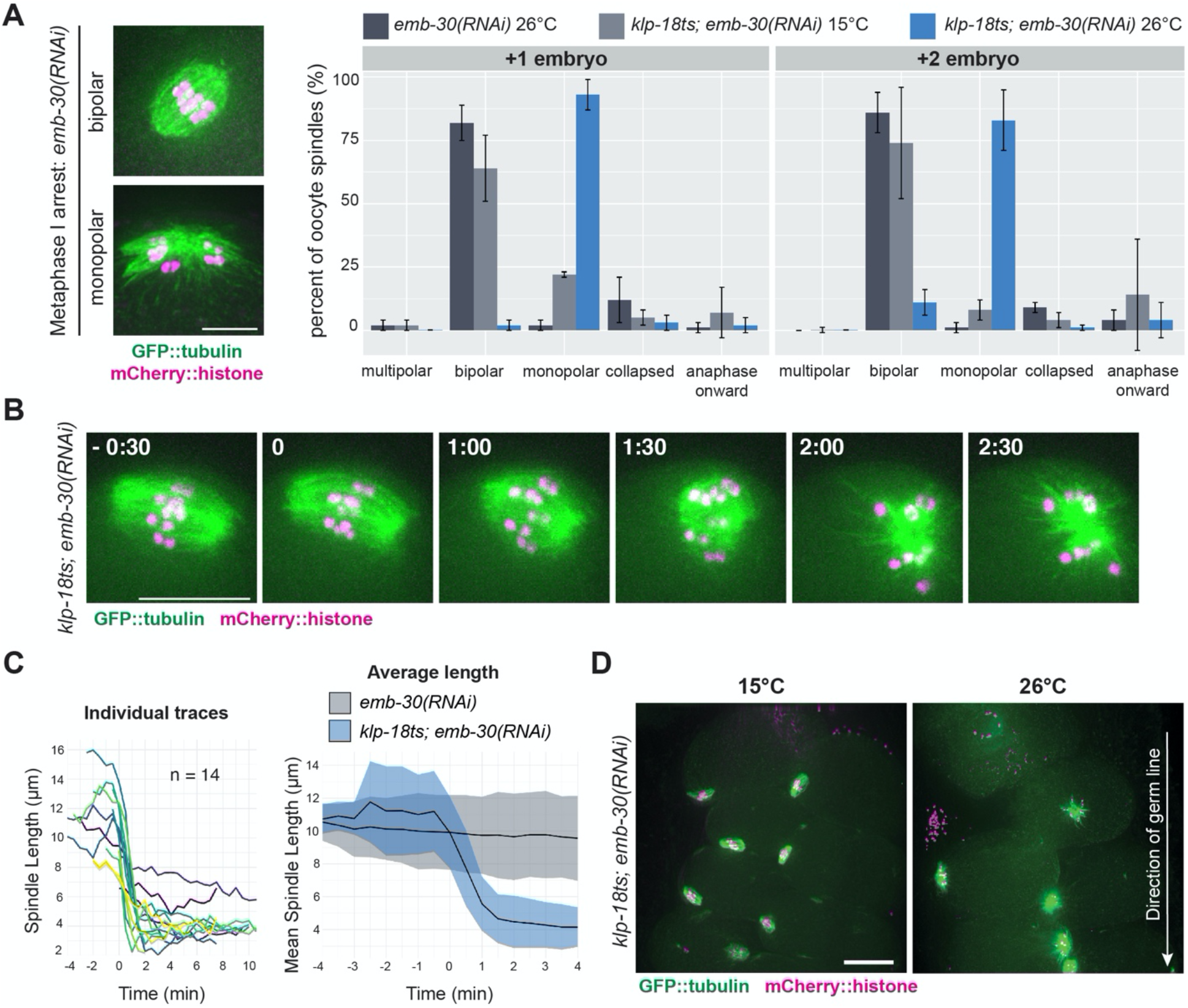
KLP-18 sorting activity is essential to maintain spindle bipolarity. a) Spindle morphology in the +1 and +2 embryos of metaphase I-arrested control *emb-30(RNAi)* (dark gray) or *klp-18ts emb-30(RNAi)* worms expressing GFP::tubulin and mCherry::histone. Spindle morphology in the *klp-18ts* strain was quantified at permissive (15°C, light gray) or restrictive (26°C, blue) temperature; example images of the categories are shown in Figure S3A. Bars represent mean percentage +/- sd; example images shown on the left. For all conditions n = 3 experiments. Scale bar = 5μm. b) A metaphase-I arrested spindle was filmed a in *klp-18ts* worm expressing GFP::tubulin and mCherry::histone; the spindle shortens and then forms a monopole. Timestamps represent minutes; t = 0 was set as first frame after shortening began. Scale bar = 10μm. c) Spindle length measurements for individual spindles are shown on left. Average +/- sd for control *emb-30(RNAi)* (n = 7, gray) and *klp-18ts emb-30(RNAi)* (n = 14, blue) are shown on right, average is the black line and sd is shown as shading. For *klp-18ts emb-30(RNAi)*, time = 0 min was set as first frame after shortening began, for control *emb-30(RNAi)*, time = 0 was set as first frame acquired. Scale bar = 10μm. d) Representative images of *klp-18(or447ts); emb-30(RNAi)* germ lines at 15°C and 26°C. Arrow indicates direction of germ line; the most recently arrested spindle is at the top of each image. Scale bar = 15μm.

Next, we used live imaging to confirm these results and to visualize the dynamics of spindle reorganization. Upon dissecting and mounting *klp-18ts; emb-30(RNAi)* embryos at room temperature (23-25°C), previously assembled bipolar spindles shortened and then formed a monopole within 2 minutes (Fig 4B and 4C, Movie S1), confirming that KLP-18 activity is essential to stabilize spindles after bipolarity is achieved. We again confirmed persistent KLP-18 and MESP-1 localization to the monopolar spindle in temperature-shifted metaphase-arrested oocytes (Fig S3E), suggesting that KLP-18 remains spindle-associated but is inactive. Taken together, these results show that KLP-18 activity is essential to maintain a bipolar spindle and that this activity relies on the KLP-18 C-terminal microtubule binding domain.

## DISCUSSION

This work is the first biochemical characterization of kinesin-12/KLP-18 and MESP-1 and provides the first mechanistic dissection of motor activity in a system in which kinesin-12 is naturally the dominant microtubule sorting motor. We propose a model in which KLP-18 exists in a folded autoinhibited state that is alleviated through MESP-1 binding; MESP-1 binds to a region overlapping with the hinge and disrupts the folding of the stalk to activate C-terminal microtubule binding (Fig S4B). This could allow KLP-18 to bind one microtubule with its motor domain and another at its C-terminus, enabling it to crosslink and slide microtubules within the spindle. In addition to unhinging KLP-18, we hypothesize that MESP-1 also promotes the initial targeting of the complex to spindle microtubules, as evidenced by its ability to bind microtubules *in vitro* and its localization to spindles *in vivo* in the *klp-18ts* and *klp-18ΔC-stalk* mutants (conditions in which the C-terminal binding site is inactivated or deleted). However, MESP-1 was not sufficient to localize GFP-KLP-18ΔC-stalk to microtubules *in vitro*, suggesting that multiple associations between the KLP-18/MESP-1 complex and microtubules may be required for stable spindle targeting *in vivo*. Consistent with this view, our previous studies showed that a KLP-18 mutant with the motor domain deleted, *klp-18(ok2519)*, abolished both KLP-18 and MESP-1 localization to the monopolar spindle [12], showing that the motor domain is essential for spindle localization.

MESP-1 is predicted to be a heavily disordered protein and was difficult to purify without the stabilizing MBP tag (Fig S1B and S1C). This suggests that MESP-1 may be disordered and inactive before it associates with KLP-18, which may explain why MESP-1 cannot localize to spindle microtubules when KLP-18 is depleted [12]. Our previous work also implicated MESP-1 as a functional ortholog of TPX2 in *C. elegans* [12], which in combination with our current findings stimulates additional hypotheses that could be tested in the future. First, it is possible that once MESP-1 targets KLP-18 to microtubules, MESP-1 may regulate KLP-18 motor activity through persistent association to both the microtubule and the motor (Fig S4B), similar to TPX2 regulation of kinesin-12/Kif15 [17, 19] and kinesin-5/Eg5 [40] in mammalian systems. In addition, TPX2 has been shown to inhibit Kif15 motility and help Kif15 associate more tightly to microtubules *in vitro* [16, 18, 19]; it remains to be seen if MESP-1 regulates KLP-18 in a similar way. Lastly, the relatively weak interaction between KLP-18 N-stalk and MESP-1 (Fig 3A) that appears to be stimulated by the addition of microtubules (Fig 3D and E) suggests that the interaction between MESP-1 and KLP-18 may require microtubules as has been shown for *Xenopus* and mammalian TPX2 and kinesin-12 [16, 28].

Once associated with spindle microtubules, KLP-18 is unable to maintain spindle bipolarity if the C-terminal microtubule binding domain is disrupted, even if the complex is still associated with microtubules (Fig S4C). Although we were not able to explicitly measure the force generated by this motor, the lack of microtubule sorting, evidenced by the monopolar spindle phenotype observed in three *klp-18* mutants, suggests that a direct stalk-microtubule interaction facilitates force generation in the meiotic spindle. Notably, recent photobleaching experiments revealed rapid poleward flux of microtubules within oocyte metaphase spindles *in vivo* [41], suggesting that spindle maintenance requires constant outward force on microtubules that may be provided by KLP-18. However, future biophysical studies are needed to confirm this model.

In mammalian somatic cells, kinesin-5 generates outward force on microtubules to achieve spindle bipolarity [42–44], and when kinesin-5 function is altered, kinesin-12 can take over this role [13–15, 45, 46]. The molecular basis for this activity has been characterized *in vitro* and in cell culture [16–21, 47]. Our work confirms that certain properties of kinesin-12s are shared across species: presence of discrete coiled-coil domains in the C-terminal stalk, the requirement of a C-terminal non-motor microtubule binding site, self-inhibition of non-motor microtubule binding, and regulation by a TPX2-like adaptor protein. Further characterization of the mechanism of KLP-18 self-inhibition, activation by MESP-1, and force generation on microtubules will be essential to fully understand the role of KLP-18 in meiosis and to put KLP-18 in proper context with other essential motors, particularly mammalian kinesin-12 and kinesin-5.

Previously, we and others have proposed that KLP-18 acts to organize microtubules and provide outward force on the spindle during anaphase [48, 49], but this hypothesis has not been directly tested. Understanding the role of KLP-18 may help to resolve different models of chromosome segregation proposed in *C. elegans* oocytes (reviewed in [1, 50]) [48, 49, 51–57]. Further characterization of KLP-18’s role in spindle assembly, spindle maintenance, and chromosome segregation will be essential to fully understand *C. elegans* oocyte meiosis.

## MATERIALS AND METHODS

### C. elegans Strains

OD868: *ltSi220[pOD1249/pSW077; Pmex-5::GFP::tbb-2::operon_linker::mCherry::his-11; cb-unc-119(+)] I* [58]

HR1160: *klp-18(or447ts), dpy-20 IV* (gift from Bruce Bowerman)

SMW29 (Δhinge): *klp-18(wig2) IV; unc-119(ed3) III; ltIs37[pAA64; pie-1::mCherry::his-58; unc-119(+)] IV; ltIs25[pAZ132; pie-1::GFP::tba-2; unc-119 (+)] IV* (transgenes were from strain OD57, gift from Arshad Desai)

SMW34 (ΔC-stalk): *klp-18(wig3), ieSi38 [sun-1p::TIR1::mRuby::sun-1 3’UTR + Cbr-unc-119(+)] IV; unc-119(ed3) III; mMaple3::tba-1 I*

SMW36: HR1160 x OD868 (*klp-18(or447ts), dpy-20 IV; ItSi220[pOD1249/pSW077; Pmex-*

*5::GFP::tbb-2::operon_linker::mCherry::his-11; cb-unc-119(+)] I)*

### Generation of *Δhinge* and *ΔC-stalk klp-18* mutant strains

The endogenous *klp-18* gene was edited using a CRISPR-Cas9 based approaches similar to what has previously been described [48, 59–61]. Genomic DNA was cut through crRNA targeting to two cut sites: one at the 5’ end of the desired mutation and one at the 3’ end. A ssDNA repair template homologous to sequences upstream and downstream of the two cut sites was used to bridge the excised DNA. For ΔC-stalk (SMW34), residues 635 to 932 of KLP-18 were deleted. For Δhinge, residues 557-770 were deleted. The *dpy-10* co-injection marker was used to screen successful genome editing events. See below for a table for sequence and concentration of all oligonucleotides used for CRISPR-Cas9 directed cutting and homology directed repair. First, 13.6µM Alt-R CRISPR-Cas9 tracrRNA (IDT), *dpy-10* crRNA, and both gene specific crRNA guides were incubated at 95°C for 5min then 10°C for 5min in a thermocycler. Half of this mixture was removed and placed into a cold PCR tube on ice, then 27µM recombinant Cas9 (IDT) was added and incubated at room temperature for 5min. The *dpy-10* ssDNA oligo (IDT) and gene specific ssDNA ultramer (IDT) repair templates were then added and nuclease free water was added to a final injection mix volume of 5µL. The injection mix was loaded into a capillary tube through a mouth pipette then injected into the germ line of young adult worms. Injected P0s were recovered in M9, then pooled onto a single plate and allowed to recover at 15°C overnight. The genetic background of each mutant can be found in the “*C. elegans* Strains” section above. The next day, injected P0s were singled out to individual plates, grown at 20°C, and allowed to lay eggs. The resulting F1 progeny were screened for rollers or dumpys indicating *dpy-10/+* or *dpy-10/dpy-10*, respectively [60]. Individual animals from “jackpot” broods (plates with many dumpys and rollers) were isolated, allowed to produce progeny, then screened with single worm PCR. Mutations in both strains were confirmed through Sanger sequencing. The primers used to screen and sequence mutants are shown in the table below. Both the Δhinge and ΔC-stalk mutations are embryonic lethal (Figure 2A), therefore these strains were propagated through single worm PCR screening of isolated adults in each generation. Progeny of adults heterozygous for the mutations were used for experiments, with the assumption that ¼ will be homozygous wild-type, ½ will be heterozygous, and ¼ will be homozygous mutant.

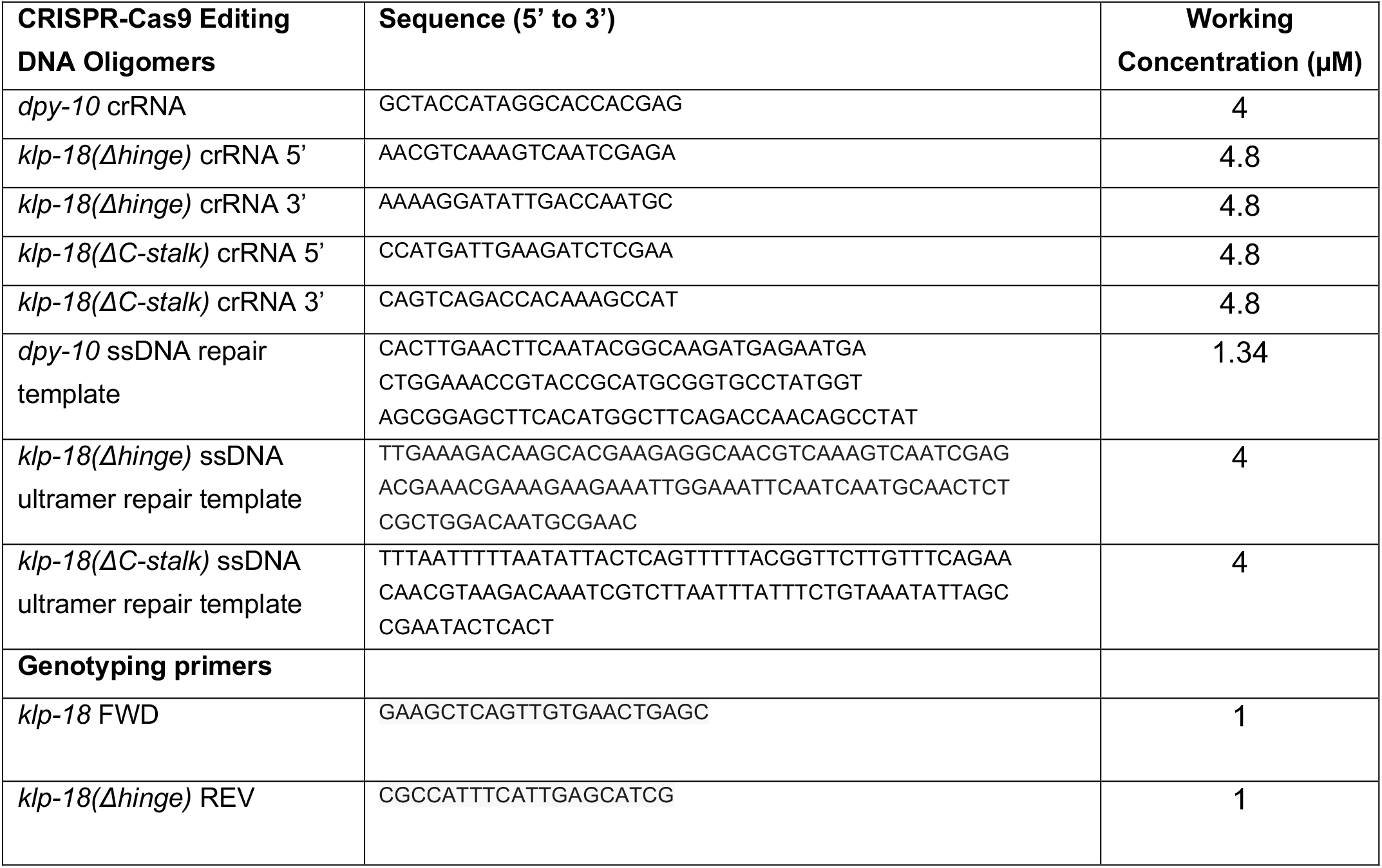

### Protein Domain analysis

KLP-18 coiled-coil domains were identified using Paircoil2 coiled-coil prediction software (http://cb.csail.mit.edu/cb/paircoil2/) [62]. Probability scores were calculated with a cutoff of 0.5. MESP-1 disordered regions predicted using PONDR (http://www.pondr.com/) [63].

### Protein Expression and Purification

Purification results, affinity tag, and expression system for each protein can be found in Figure S1A. *C. elegans* cDNA was amplified from extracted mRNA from wild type worms using the iScript Select cDNA Synthesis kit (Bio-Rad). KLP-18 cDNA was amplified from whole worm cDNA with gene specific primers and Q5 DNA polymerase, then used for cloning all KLP-18 pET expression constructs. Δhinge was created via site-directed mutagenesis (NEB) from the stalk construct. MESP-1 cDNA was amplified from a GST-MESP-1 construct [12] and inserted into an MBP vector (gift of Laura Lackner and Marijn Ford). pET His6 GFP TEV LIC cloning vector (1GFP [64]) was a gift from Scott Gradia (Addgene plasmid # 29663; http://n2t.net/addgene:29663 ; RRID:Addgene_29663). KLP-18 cDNA was inserted into the GFP construct via Gibson Assembly to create the GFP-stalk construct, and the GFP-KLP-18ts and GFP-ΔC-stalk were created via site-directed mutagenesis (NEB) from this GFP-stalk construct. KLP-18 cDNA was inserted into the MBP construct via restriction digest to make MBP-stalk.

All proteins were purified using the same protocol and the same buffers. All expression vectors were transformed into BL21 DE3 *E. coli* cells and grown at 37°C to an O.D. of ∼0.6. Cells were induced with 0.1mM IPTG and grown for varying expression times and at varying temperatures; see Figure S1A for growth conditions for individual proteins. Growth cultures were spun at 4700rpm and resuspended in lysis buffer (80mM PIPES pH 6.8, 2mM MgCl_2_, 1mM EGTA, 250mM NaCl, 5-10% glycerol, 0.02% Tween, Leupeptin, Aprotinin, Pepstatin, 2mM imidazole) then cells were lysed with 1mg/mL lysozyme (incubated for 20min at 4°C) and with sonication. Lysate was cleared by centrifugation for 45min at 11,900rpm in a Ti50.2 rotor. Ni-NTA resin was equilibrated with lysis buffer then added to cleared lysate and incubated at 4°C for 1-2 hours. Slurry was applied to a plastic column and washed with 30mL lysis buffer + 20mM imidazole. Bound protein was eluted with 5-10mL of lysis buffer + 500mM imidazole. Ni-NTA elution was either kept at 4°C overnight before application or applied directly to HiLoad 26/600 Superdex 200 gel filtration column run on Aktaprime FPLC system (GE Biosciences) and equilibrated with lysis buffer. Eluted fractions from individual peaks were tested by SDS PAGE and fractions containing pure protein of interest were combined, concentrated, frozen by dripping into liquid N_2_, and stored at -80°C.

### Microtubule co-sedimentation assay

100µM purified porcine brain tubulin (self-purified, gift of Sarah Rice) was incubated with 1mM DTT and 1mM GTP in BRB80 (80mM PIPES pH 6.8, 2mM MgCl_2_, 1mM EGTA) on ice for 5min then spun at 80K rpm for 10min at 4°C. Supernatant was removed and incubated at 37°C for 1 hour. Taxol was added stepwise to the following final concentrations at 37°C: 1µM taxol then incubated for 10min, 10µM taxol then incubated for 10min, and 100µM taxol then incubated for 15min. Polymerized microtubules were diluted 1:1 in BRB80 + 50µM taxol and kept at room temperature until use. For subtilisin treatment, 1mg/mL subtilisin was added to 20µM polymerized tubulin and incubated overnight at 37°C. Reaction was quenched with 2mM final concentration of PMSF then incubation at 37°C for 1 hour then spun over a cushion (200µL BRB80, 60% glycerol, 20µM taxol) at 80K rpm for 20min. Cushion and pellet were washed in 50µL BRB80 + 20µM taxol, then resuspended in 10µL BRB80 + 50µM taxol.

Proteins of interest were thawed from -80°C storage and pre-cleared by spinning at 80K rpm for 10 minutes at 25°C. Soluble protein in the supernatant was removed and added to 5µM microtubules quickly to maintain protein solubility. Reactions were assembled in BRB80 + 20µM taxol with equal salt, detergent, and glycerol concentrations based on the final purification buffers (see “Protein Purification” section for details) to a final volume of 25µL. Exact protein concentrations for each experiment can be found in “Figure Quantification” section of Materials and Methods. Reactions were incubated at room temperature for 30min, then spun through a 100µL BRB80 + 40% glycerol + 20µM taxol cushion at 90K rpm for 15min at 25°C. 25µL from the very top of the solution was removed and added to 2X SDS Laemmli Sample Buffer to make the “supernatant” sample. The cushion was washed with BRB80 + 20µM taxol, removed, then pellet was washed with BRB80 + 20µM taxol. Pellet was resuspended with 25µL cold BRB80 + 10mM CaCl_2_then added to 2X SDS Laemmli Sample Buffer to make the “pellet” sample. All spins performed in TLA120.2 rotor (Beckman).

Supernatant and pellet samples were probed by Western Blot using the following antibody concentrations: 1:5000 anti-6XHis-HRP (Abcam), 1:5000 mouse anti-tubulin (Invitrogen), 1:5000 anti-mouse HRP (Invitrogen). See “Figure Quantification” section of Materials and Methods for Western Blot quantification details.

### Microtubule bundling assay

Fluorescent microtubules were polymerized by incubating 20µM porcine brain tubulin and 2µM TMR-tubulin (Cytoskeleton) with 1mM DTT and 1mM GTP in BRB80 (80mM PIPES pH 6.8, 2mM MgCl_2_, 1mM EGTA) on ice for 5min and incubated at 37°C for 2min. Taxol was added stepwise to the following final concentrations at 37°C: 0.2µM taxol then incubated for 10min, 2µM taxol then incubated for 10min, and 20µM taxol incubated for 10min. Polymerized microtubules were kept at room temperature until use.

Proteins of interest were thawed from -80°C storage and pre-cleared by spinning at 80K rpm for 10 minutes at 25°C. Protein concentrations were measured by Bradford Assay, then added to 25µL reactions with 200nM TMR-microtubules in BRB80 + 20µM taxol with equal salt, detergent, and glycerol concentrations based on the final purification buffers (see “Protein Purification” section for details). Reactions were incubated at room temperature for 30min, then fixed with 1% glutaraldehyde and incubated for an additional 3min. Reactions were then squashed onto a poly-L-lysine slide, sealed with nail polish, and imaged on a Spinning Disk Confocal microscope (see “Time-lapse imaging” section for microscope details). For exact protein concentrations and quantification details, see “Figure Quantification” section of Materials and Methods.

### Embryonic lethality assay

Embryonic lethality scoring was performed as previously described [48] with slight modifications: L4 progeny of unbalanced *klp-18* mutant strains (SMW29 or SMW34) were isolated onto individual NGM plates and allowed to lay eggs at 20°C for 24 hours before being transferred to a fresh NGM plate. The eggs were allowed to hatch for 24 hours at 20°C and then the total progeny, unhatched eggs and hatched worms, were scored. This process was repeated two times per isolated worm for a total of three scored plates per worm. Following the third day of egg laying, parent worms were screened for genotype (homozygote background, *klp-18* heterozygote mutant, or *klp-18* homozygote mutant) by PCR.

### Fixed imaging

For quantification of spindle morphology, intact worms were fixed in EtOH: ∼30-45 worms were picked into a 15°C drop of M9 (22mM KH_2_PO_4_, 22mM Na_2_HPO_4_, 85mM NaCl, 1mM MgSO_4_), the drop was dried with Whatman paper, and 10µL 100% EtOH was added directly to worms. The EtOH was allowed to dry completely and another drop of 100% EtOH was added, and this was repeated for a total of 3 times. A 1:1 mixture of Vectashield:M9 was added to the worms then a coverslip was added and sealed with nail polish. Slides were stored at 4°C until imaging. Slides were visualized on a DeltaVision Core microscope (G.E.) and spindle morphology was quantified by eye with a 40X objective or by taking a snapshot of the spindle with a 100X objective. Oocyte spindles in embryos in the +1 or the +2 positions were quantified as indicated on the figure.

Immunofluorescence was performed as previously described [65, 66]. Oocytes were dissected into drops of M9, snap-frozen in liquid N_2_, freeze cracked, and fixed in -20°C MeOH for 35 minutes. Slides were washed twice with PBS, blocked with Abdil (1X PBS, 4% BSA, 0.1% Triton X-100, 0.02% Sodium Azide) for 30 minutes at room temperature, then primary antibody was applied overnight at 4°C. The next day slides were washed three times with PBST at room temperature, incubated with secondary antibody for 2 hours at room temperature, washed three times with PBST, incubated with Hoechst for 15 minutes at room temperature, washed twice with PBST, then mounted in mounting media (90% glycerol, 20mM Tris pH 8.8, 0.5% p-phenylenediamine), and sealed with nail polish. Primary antibodies and concentrations used: rat anti-KLP-18 (1:500, gift of Olaf Bossinger [9]), rabbit anti-MESP-1 (1:3000 [12]), rabbit ASPM-1 (1:5000 [12]). Secondary antibodies and concentrations used: goat anti-rat (1:500, Invitrogen), goat anti-rabbit (1:500, Invitrogen). Slides were imaged using a DeltaVision Core microscope with a 100X objective (NA = 1.40). All image acquisition and processing was performed using softWoRx software (GE Biosciences). Image stacks were acquired with 0.2µm z-steps. All immunofluorescence images are full maximum-intensity projections of the entire spindle structure and are not deconvolved unless indicated in the figure legend.

Microscopy was performed at the Biological Imaging Facility at Northwestern University.

### GFP-tagged protein recruitment to microtubules

Fluorescent microtubules were polymerized as previously described [21]. First, GMPCPP seeds were nucleated by incubating 13µL PEM104 (104mM PIPES pH 6.9, 1.3mM EGTA, 6.3mM MgCl_2_), 2.2µL 10mM GMPCPP, 2.2µL DMSO, 4µL 13mg/mL homemade porcine brain tubulin (gift of Sarah Rice), and 4µL 10mg/mL TMR-tubulin (Cytoskeleton) at 37°C for 40min. A microtubule elongation mixture of 13µL PEM104, 2.2µL 10mM GTP, 2.2µL DMSO, 2µL 13mg/mL homemade porcine brain tubulin, and 1.5µL 1mg/mL TMR-tubulin (Cytoskeleton) was incubated at 37°C for 1min, 1.5µL GMPCPP seed mixture was added, then reaction was incubated at 37°C for 40min. Microtubules were stabilized by adding 2µL of stabilization buffer (38.6µL BRB80, 0.5µL 100mM GTP, and 6µL 2mM taxol dissolved in DMSO) then kept at room temperature until use. Microtubules were used within one week of polymerization. Proteins of interest were thawed from -80°C storage then added to 25µL reactions with 5µL of 1:20 diluted TMR-microtubules in BRB80 + 20µM taxol with equal salt, detergent, and glycerol concentrations based on the final purification buffers (see “Protein Purification” section for details). Reactions were incubated, fixed, and imaged exactly as described in “microtubule bundling assay” above.

### MBP pulldown

1µM N-stalk / C-stalk and 2µM MBP / MBP-MESP-1 were added to a 100µL reaction in BRB80 (80mM PIPES pH 6.8, 2mM MgCl_2_, 1mM EGTA) with equal salt, detergent, and glycerol concentrations based on the final purification buffers (see “Protein Purification” section for details). Equal volume of protein was added to 200µL 1X Laemmli Sample Buffer for the “input” gel samples. Reactions were incubated at 4°C overnight, then equilibrated amylose resin (New England Biolabs) was added to pull out MBP-tagged proteins. Amylose resin was incubated with reactions for 1 hour at 4°C, then resin was washed 4 times with 900µL BRB80. 300µL of final wash was saved for acetone precipitation to make “wash” samples. To elute bound proteins from amylose resin, 900µL BRB80 + 50mM NaCl + 10mM maltose was added to beads and incubated for 20min at 4°C. 300µL of eluate was removed for acetone precipitation to make “elution” gel samples.

Wash and elution samples were concentrated via acetone precipitation as follows: 1.2mL of - 20°C acetone was added to 300µL wash or elution samples, vortexed, and incubated at -20°C overnight. The next day samples were spun at 16K x g for 10min at room temperature and incubated at 50-55°C for ∼20min until the acetone was completely evaporated and the pellets were dry. Pellet was resuspended in 40µL 1X Laemmli Sample Buffer and boiled for 10min at 95°C to make gel samples. Samples were run on a Western blot and probed with 1:5000 anti-6XHis-HRP to visualize His-tagged KLP-18 truncations. To visualize MBP or MBP-MESP-1 samples were run on an SDS-PAGE gel and stained with Coomassie. Input lane represents 1% of protein added to the reaction, and wash/elution lanes represents 12.5% of all protein in those samples.

### RNA interference

Individual RNAi clones picked from an RNAi feeding library [67, 68] were used to inoculate Luria broth (LB) plus ampicillin (100 μg/mL) and grown overnight at 37°C. These cultures were used to seed nematode growth medium (NGM) / ampicillin (100 μg/mL) / 1 mM IPTG plates, and the plates were left overnight at room temperature. Synchronized L1 worms were plated on induced plates and grown at 15°C for 5-6 days until they became gravid adults.

### Time-lapse imaging

Worms were prepared for two-color live imaging as previously described [69]. *klp-18ts; emb-30(RNAi)* and control *emb-30(RNAi)* adult worms were kept at 15°C and room temperature prior to dissection, respectively. Oocytes were dissected out of worms in a drop of room temperature L-15 Blastomere buffer (final concentrations in ddH_2_O: 60% Leibovitz L-15, 20% heat inactivated FBS, 25mM HEPES pH 7.5, 0.5mg/mL Inulin) then a coverslip was added. Oocytes that settled onto the coverslip were filmed. All live imaging was performed at ambient temperature (∼23-25°C).

Imaging was performed using a spinning-disk confocal microscope with a 63X HC PL APO 1.40 NA objective lens. A spinning-disk confocal unit (CSU-X1; Yokogawa Electric Corporation) attached to an inverted microscope (Leica DMI6000 SD) and a Spectral Applied Imaging laser merge ILE3030 and a back-thinned electron multiplying charge-coupled device camera (Evolve 521 Delta) were used for image acquisition. The microscope and attached devices were controlled using MetaMorph Image Series Environment software (Molecular Devices). Frames were acquired every 30 seconds at 2µm z-stack intervals.

### Hydrodynamic Analysis

A purification of MBP-Stalk was split and diluted into BRB80 + 300mM NaCl (high salt) or BRB80 + 20mM NaCl (low salt). The high salt and low salt dilutions were then applied to HiLoad 26/600 Superdex 200 column equilibrated with appropriate buffer. Identical fractions were collected for each dilution and probed by Western Blot using rabbit anti-KLP-18 at 1:5000 and anti-rabbit-HRP at 1:5000. See “Figure Quantification” section for further details.

### Western Blots

For *klp-18(or447ts)* whole worm Western blots, control (OD868) or *klp-18(or447ts)* (SMW36) plates were shifted to 26°C for 1 hour. 50-100 worms were picked from shifted plate to pre-warmed unseeded plate for 5min to avoid transfer of bacteria, then washed off plate with room temperature M9. Worms were pelleted by spinning at 800xg for 1min, supernatant was removed, and an equal volume of 2X SDS Laemmli Sample Buffer was added. Gel samples were boiled for 10min at 95°C, briefly vortexed, then boiled for an additional 10min at 95°C. Volume of sample corresponding to 50 worms was loaded onto gel. Antibodies used: rat anti-KLP-18 1:1000, anti-rat-HRP 1:5000 (Invitrogen), mouse anti-tubulin 1:5000, anti-mouse-HRP 1:5000 (Invitrogen).

Samples were run on 8-12% SDS-PAGE gel and transferred to a nitrocellulose membrane using a Trans-Blot Turbo Transfer System (BioRad). Membrane was blocked in 5% milk in TBS + 0.1% Tween blocking solution, incubated with primary antibody in blocking solution at room temperature for 1 hour or at 4°C overnight, washed in TBST, incubated in secondary antibody at room temperature for 1 hour, washed in TBST, incubated with Clarity Western ECL substrate (BioRad) for 2 minutes, then imaged.

### Figure Quantification

*Figure 1B:* Western blots were developed on film then scanned, and the ImageJ “analyze gels” function was used to quantify band intensity in each lane [70, 71]. For each reaction, the total band intensity was calculated by adding the “supernatant” and “pellet” band intensities, and “shift to pellet (%)” was calculated by dividing the “pellet” band intensity by the total band intensity for the reaction. The mean +/- sd for n = 3 independent experiments are shown. For each experiment quantified, [tubulin] = 5µM, [N-stalk] = 1.0µM-1.26µM, [C-stalk] = 0.14µM-1µM. In the representative blots shown for 1B, left: [N-stalk] = 1.26µM and [C-stalk] = 0.53µM. For the subtilisin experiment shown in 1B, right: [C-stalk] = 0.14µM for all reactions. For non-subtilisin experiment: Shift to pellet (%) +/- sd for N-stalk (-MT) = 4.5% +/- 4.9%, N-stalk (+ MT) = 3.2% +/- 3.6%, C-stalk (-MT) = 1.6% +/- 2.8%, C-stalk (+ MT) = 52.7% +/- 9.7%. For subtilisin experiment: Shift to pellet (%) +/- sd for C-stalk (-MT) = 6.0 +/- 6.7, C-stalk (+ MT) = 50.8% +/- 7.0%, C-stalk (+ sMT) = 26.7% +/- 7.0%.

*Figure 1D:* Microtubule bundling assays were imaged on a spinning disk microscope with consistent exposure times across the experiment. Images were quantified in ImageJ: raw images were made into a grayscale composite then an automatic threshold using the “Threshold” command was applied. Across an experiment, a threshold was applied to each image individually but with the same automatic thresholding algorithm. After the thresholding was applied, fluorescent particles were selected and the “mean size” was calculated. This measurement is the mean size of all fluorescent particles in an image and represents the degree of microtubule bundling (large bundles will have a larger mean size than individual microtubules). To normalize mean bundle area, the area from experimental images was divided by the mean area of all of the buffer-only control images, making the mean size of unbundled microtubules equal to 1. Each plotted data point is one image’s normalized bundle area. Two experiments are shown on the graph and the same result (Δhinge bundles MTs while full length stalk does not) was shown in 6 total experiments. In each reaction, [tubulin] = 200nM. Mean “Normalized bundle area (A.U.)” and n for each condition are buffer alone: 1, n = 62; 1µM stalk: 1.22, n = 67; 2µM stalk: 1.59, n = 62; 1µM Δhinge: 2.63, n = 63; 2µM Δhinge: 5.27, n = 62. Using a two-sample equal variance Student’s t-test, all conditions compared to buffer and Δhinge compared to equal concentrations of stalk showed statistically significant (p< 0.01) increases in normalized bundle area.

*Figure 2B:* Spindle morphology in control or *klp-18(or447ts)* worms fixed in EtOH was quantified by eye using a 40X objective or by taking snapshots at 100X. The percentages shown in Figure 2B are calculated by considering only pre-anaphase oocyte spindles (multipolar, bipolar, monopolar); representative images are shown. However, we did assess all +1 spindles (including anaphase and MII spindles), and that full dataset is graphed in Figure S3A. For *klp-18(or447ts)*, 4 experiments were quantified (for 15°C n = 28, 16, 32, 29 pre-anaphase spindles; for 26°C n = 27, 21, 42, 42 pre-anaphase spindles); for control 26°C, 3 experiments were quantified (n = 12, 11, 14 pre-anaphase spindles). Mean “percent of pre-anaphase +1 oocytes (%)” +/- sd for control 26°C: multipolar = 6% +/- 10%, bipolar = 94% +/- 10%, monopolar = 0% +/- 0%. Mean “percent of pre-anaphase +1 oocytes (%)” +/- sd for *klp-18(or447ts)* 15°C: multipolar = 10% +/- 7%, bipolar = 67% +/- 7%, monopolar = 23% +/- 10%. Mean “percent of pre-anaphase +1 oocytes (%)” +/- sd for *klp-18(or447ts)* 26°C: multipolar = 1% +/- 1%, bipolar = 4% +/- 4%, monopolar = 95% +/- 5.

*Figure 2C and D:* Fluorescence intensity of KLP-18 or MESP-1 immunofluorescence staining at poles of fixed oocyte spindles. MI or MII bipolar or monopolar spindles were quantified for each condition. Raw undeconvolved images were used for quantification and each image was acquired at similar exposures and laser power. For the *klp-18ΔC-stalk* and *klp-18Δhinge* strains, each slide contained a mix of bipolar and multipolar spindles allowing for an internal control. Every image of a spindle on the same slide was acquired identically. All image analysis was performed in ImageJ [70, 71]. First, each pole of the bipolar spindle or single monopole was qualitatively identified in the tubulin channel. A 5-slice sum projection was then made containing 2 z-stacks above and below the center z-slice at the center of the pole for each channel, then a ROI was drawn around the pole in the tubulin channel of the sum projection. The exact ROI was applied to the KLP-18 and MESP-1 channels, and the integrated density (mean pixel intensity x area) was recorded for each channel. To calculate background, the same ROI was moved to an area of the cytoplasm adjacent to each pole and the same intensity measurements were recorded. The background measurements were then subtracted from the pole measurements. For bipolar spindles, the two background-subtracted poles were averaged together. To calculate final fluorescence intensity on spindle poles, the background-subtracted KLP-18 or MESP-1 intensity was divided by the background-subtracted tubulin intensity to normalize for variability in staining or the amount of microtubules present on mutant spindles. Each data point represents one spindle. For *klp-18ts* bipolar n = 21 spindles; *klp-18ts* monopolar n = 38 spindles; *klp-18ΔC-stalk* bipolar n = 38 spindles; *klp-18ΔC-stalk* monopolar n = 18 spindles; *klp-18Δhinge* bipolar n = 23 spindles; *klp-18ΔC-stalk* monopolar n = 22 spindles.

*Figure 3B:* Quantified exactly as described in Figure 1B, except some blots were imaged with an Azure Biosystems digital imager. Concentrations for each experiment quantified: [tubulin] = 5µM, [MBP-MESP-1] = 1.0-5.77µM. In the representative blots shown, [MBP-MESP-1] = ∼1.0µM (non-subtilisin experiment) and 5.77µM (subtilisin experiment). Two non-subtilisin experiments were quantified and three subtilisin experiments were quantified. Shift to pellet (%) +/- s.d for MBP-MESP-1: (-MT) = 2.9% +/- 4.1%, (+ MT) = 56.1% +/- 3.4%. For subtilisin experiments, shift to pellet (%) +/- sd for MBP-MESP-1: (-MT) = 0.6% +/- 0.1%, (+ MT) = 41.6% +/- 10.9%, (+ sMT) = 4.5% +/- 6.4%.

*Figure 3D and E:* Images were acquired and threshold applied as described for Figure 1D. Exposure times were held constant across the experiment. To quantify GFP localization, fluorescent particles were selected in the microtubule channel then the mean intensity in the GFP channel was measured within the selection. Therefore, only GFP signal that was overlaid on microtubule bundles was measured. Mean intensity in the tubulin channel was also measured. To normalize to background, the selection was inverted to select the entire image that was not previously selected as microtubules and the mean intensity in both the GFP and red tubulin channel was measured. The mean background intensity was subtracted from mean GFP intensity for each image. ‘Normalized GFP enrichment’ was calculated by dividing background-subtracted GFP signal by background-subtracted tubulin signal to normalize for variability in the amount and size of microtubule bundles. The mean ‘Normalized GFP enrichment’ value for an experiment was calculated and averaged over n = 3 experiments. The mean +/- sd for each condition are as follows: GFP + MBP-MESP-1 = 0.14 +/- 0.1; GFP-ΔC- stalk = 0.09 +/- 0.01; GFP-ΔC-stalk + MBP-MESP-1 = 0.13 +/- 0.02; GFP-*klp-18ts* = 0.24 +/- 0.08; GFP-*klp-18ts* + MBP-MESP-1 = 0.37 +/- 0.06; GFP-stalk = 0.10 +/- 0.03; GFP-stalk + MBP-MESP-1 = 0.71 +/- 0.10. Each reaction contained 250nM GFP-tagged proteins and 1µM MBP-MESP-1 (if added as indicated).

*Figure 4A:* Quantified as in Figure 2B and S3A. For all strains and conditions, 3 experiments were quantified (control 26°C n = 52, 26, 28 +1 spindles, n = 50, 26, 27 +2 spindles; *klp-18(or447ts)* 15°C n = 42, 81, 52 +1 spindles, n = 38, 77, 55 +2 spindles; *klp-18(or447ts)* 26°C n = 39, 50, 56 +1 spindles, n = 36, 42, 56 +2 spindles). The mean percent of oocytes (%) +/- sd for each condition was calculated. For control *emb-30(RNAi)* 26°C +1 oocytes: multipolar = 2% +/- 2%; bipolar = 82% +/- 7%; monopolar = 2% +/- 2%; collapsed = 12% +/- 9%; anaphase onward = 1% +/- 2%. For control *emb-30(RNAi)* 26°C +2 oocytes: multipolar = 0% +/- 0%; bipolar = 86% +/- 8%; monopolar = 1% +/- 2%; collapsed = 9% +/- 2%; anaphase onward = 4% +/- 4%. For *klp-18(or447ts) emb-30(RNAi)* 15°C +1 oocytes: multipolar = 2% +/- 2%; bipolar = 64% +/- 13%; monopolar = 22% +/- 1%; collapsed = 5% +/- 3%; anaphase onward = 7% +/- 1%. For *klp-18(or447ts) emb-30(RNAi)* 15°C +2 oocytes: multipolar = 0%; bipolar = 74% +/- 22%, monopolar = 8% +/- 4%; collapsed = 4% +/- 3%; anaphase onward = 4% +/- 7%. For *klp-18(or447ts) emb-30(RNAi)* 26°C +1 oocytes: multipolar = 0% +/- 0%; bipolar = 2% +/- 2%; monopolar = 93% +/- 6%; collapsed = 3% +/- 3%; anaphase onward = 2% +/- 3%. For *klp-18(or447ts) emb-30(RNAi)* 26°C +2 oocytes: multipolar = 0% +/- 0%; bipolar = 11% +/- 5%; monopolar = 83% +/- 12%; collapsed = 1% +/- 1%; anaphase onward = 4% +/- 7%.

*Figure 4C:* To quantify spindle length in time-lapse movies, raw data was loaded in ImageJ and the distance between poles was measured with the line tool: spindle poles were identified in the tubulin channel by a bright circular area of tubulin, and a straight line was drawn between the outer edges of pole tubulin signal to measure the spindle length. This was done for each frame of the movie. Because spindle collapse in *klp-18ts; emb-30(RNAi)* oocytes happened at slightly different times after beginning filming, the time scale was normalized such that the first frame showing shortening was set as time = 0. Each trace for 14 spindles is shown on the left, and the mean spindle length (+/- sd) over time for the same data set is shown on the right (blue shading). For control *emb-30(RNAi)* spindles, because there was no obvious spindle collapse, the beginning of the video was set at time = -4 and the time was not re-adjusted.

*Figure S2C:* The first 11 fractions after the HiLoad 26/600 Superdex 200 void volume (fractions 43-54) were collected and run on a Western blot and developed film was scanned using a printer scanner. To calculate “% of total band intensity” the intensity of each band from fractions 43-49 was quantified. These intensities were then summed to calculate the total band intensity, and each individual fraction band intensity was divided by the total band intensity. Fractions 49-54 were not included in this calculation because they were the main elution peak of the protein, and we were interested in the fraction of protein shifted to a lower elution volume.

*Figure S3A:* Spindle morphology in *klp-18(or447ts)* worms fixed in EtOH was quantified by eye using a 40X objective or by taking snapshots at 100X. Representative images are shown. For *klp-18(or447ts)* experiments, 4 experiments were quantified (for 15°C n = 72, 54, 68, 65 spindles; for 26°C n = 59, 58, 87, 68 spindles); for control 26°C, 3 experiments (n = 57, 38, 46 spindles) were quantified. The mean percent of oocytes (%) +/- sd for each condition was calculated. For control 26°C: multipolar = 1% +/- 2%; bipolar = 26% +/- 7%; monopolar = 0% +/- 0%; collapsed = 7% +/- 4%, anaphase onward = 67% +/- 8%. For *klp-18(or447ts)* 15°C: multipolar = 4% +/- 3%; bipolar = 27% +/- 7%; monopolar = 9% +/- 4%; collapsed = 13% +/- 4%; anaphase onward = 47% +/- 6%. For *klp-18(or447ts)* 26°C: multipolar = 0 +/- 1%; bipolar = 2% +/- 2%; monopolar = 45% +/- 1%; collapsed = 22% +/- 5%; anaphase onward = 30% +/- 14%.

*Figure S3C:* Western blot band intensities were quantified using Image J. ‘KLP-18 / tubulin intensity’ was calculated by dividing KLP-18 band intensity by the tubulin band intensity in the same lane. Mean intensity +/- sd for 3 individual experiments: control 26°C = 2.06 +/- 0.78; *klp-18(or447ts)* 15°C = 2.05 +/- 0.30; *klp-18(or447ts)* 26°C = 2.69 +/- 0.85. P-value from paired one-tail Student’s t-Test comparing control 26°C to *klp-18(or447ts)* 15°C = 0.49 and control 26°C to *klp-18(or447ts)* 26°C = 0.24.

*Figure S3E:* KLP-18 and MESP-1 enrichment at pole of monopolar spindles was quantified in immunofluorescence images that exhibited characteristic traits of monopolar spindles: chromosomes out in a rosette and clearly associated to microtubule bundles. Both MI and MII spindles were quantified. First, in ImageJ, the z-slice that contained the center of the pole was qualitatively chosen in the tubulin channel and a region of interest (ROI) was drawn around the pole. The ROI was then applied to the KLP-18 and MESP-1 channel at the same z-slice, and the sum of pixel intensity was measured. A background ROI of equal size was measured in the cytoplasm for both channels at the same z-slice. To calculate enrichment of KLP-18 and MESP-1 to the pole, the pole intensity was divided by the background intensity. If this ratio was greater than or equal to 1.5 (a 50% increase), KLP-18 or MESP-1 was considered enriched. This quantification is reported on the figure.

## ACKNOWLEDGMENTS

We thank members of the Wignall lab and the WiLa ICB for support and thoughtful discussions; Gabriel Cavin-Meza, Hannah Horton, Rachel Kadzik, and Tim Mullen for critical reading of the manuscript; Sarah Rice for reagents and technical advice; Olaf Bossinger, Bruce Bowerman, Arshad Desai, Marijn Ford, Scott Gradia, and Laura Lackner for reagents; and Vladimir Gelfand, Laura Lackner, and John Marko for helpful suggestions. Microscopy was performed at the Biological Imaging Facility at Northwestern University, supported by the Chemistry for Life Processes Institute, the NU Office for Research, and the Rice Foundation. This work was supported by NIH R01GM124354 (to S.M.W.), American Heart Association Predoctoral Fellowship 17PRE33440016 (to I.D.W.), and NIH/NIGMS Molecular Biophysics Training Grant T32GM008382 (to I.D.W.).

---

**Figure S1:**
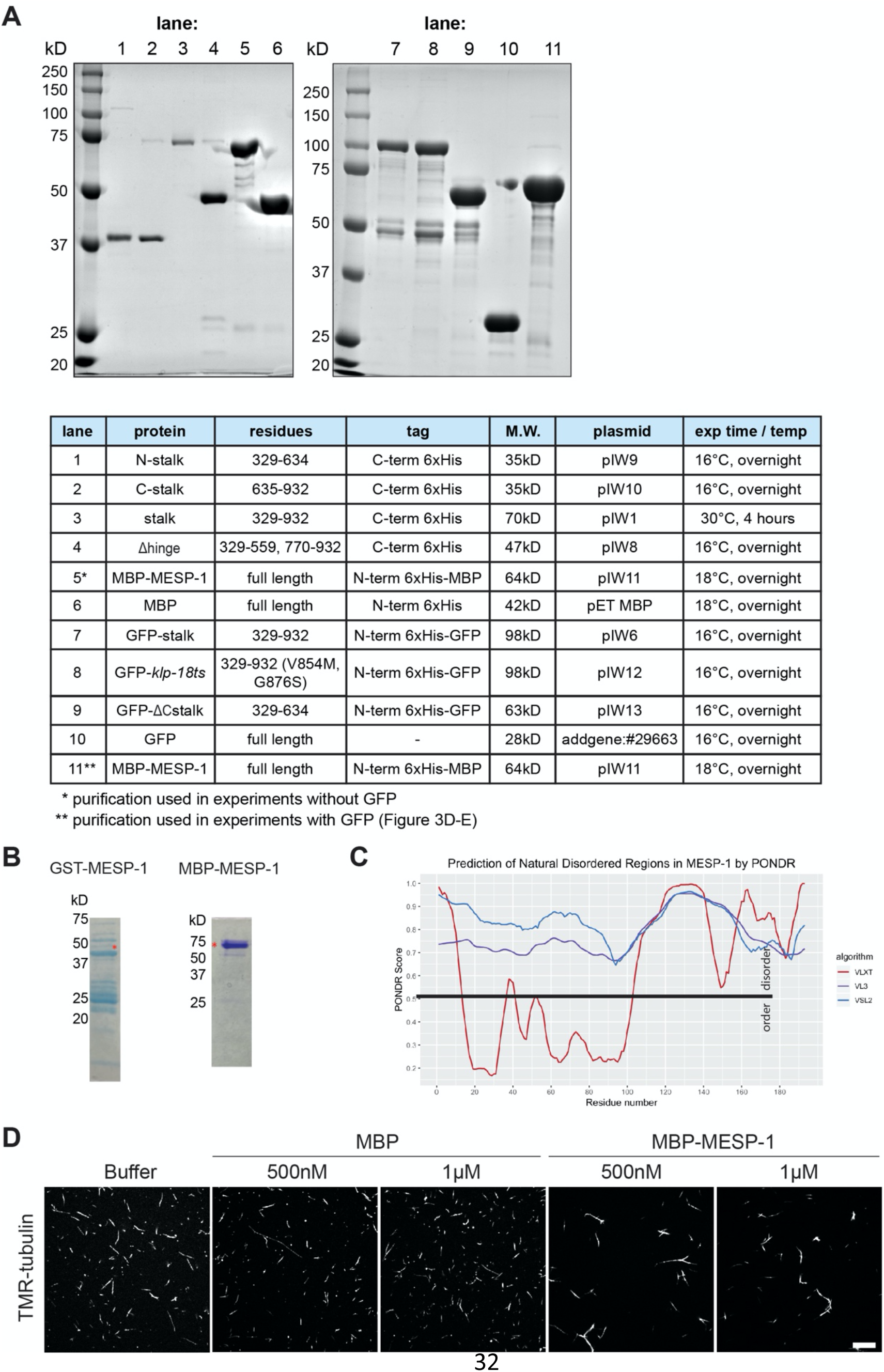
Protein expression and purification details. a) SDS-PAGE gels stained with Coomassie showing purifications of all proteins used. For purification details, see Materials and Methods. Protein of interest is major band in each lane. Each purification was confirmed by Western Blot probed with an antibody against protein of interest. The corresponding residues, affinity tag(s), molecular weight (M.W.), plasmid name, and expression time / temperature (after 0.1mM IPTG induction) for each protein are shown in the chart. The purification of MBP-MESP-1 shown in lane 5 was used in Figures S1D, 3A, 3B, and the purification shown in lane 11 was used in the GFP experiments in Figure 3D and 3E; the conditions for these purifications were the same but they were performed on different days. b) Representative purifications shown for GST- and MBP-tagged MESP-1. GST-MESP-1 and MBP-MESP-1 band marked by red asterisk. GST-MESP-1 shows more degradation during purification than MBP-MESP-1. c) Prediction of disordered regions within MESP-1 using PONDR [63]. PONDR score for three algorithms (VXLT (red), VL3 (purple), and VSL2 (blue)) shown for each residue. Residues with PONDR scores above the black line (> 0.5) are predicted to be disordered. d) MBP-MESP-1 microtubule binding activity tested by microtubule bundling assay. Representative images of TMR-microtubules incubated with buffer alone, MBP, and MBP-MESP-1. Scale bar = 10μm.

**Figure S2:**
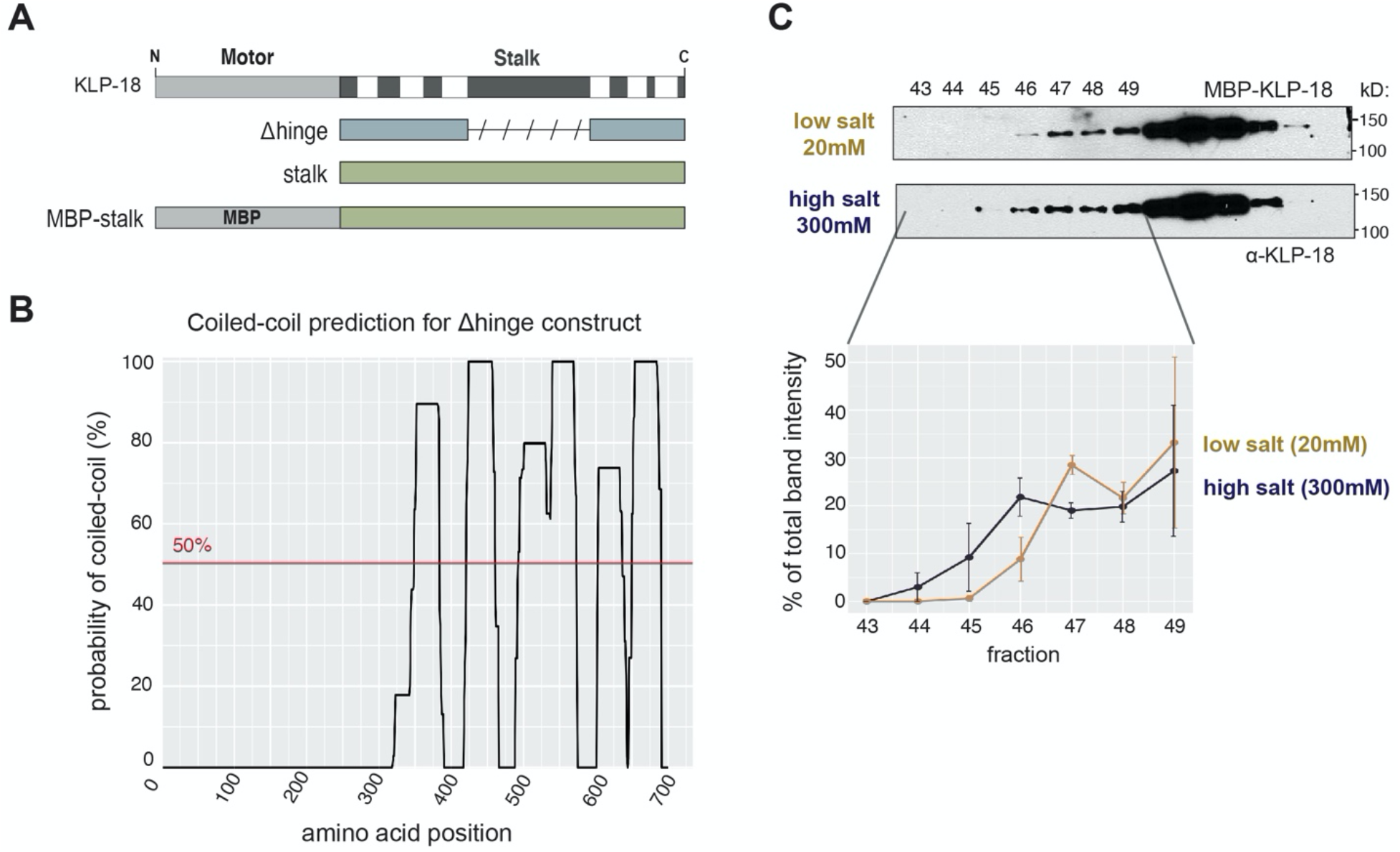
KLP-18 stalk contains a putative hinge region. a) Schematics of Δhinge, full length stalk, and MBP-stalk constructs. Slanted dashed lines show area deleted in Δhinge. b) Paircoil2 prediction for Δhinge construct with putative hinge deleted. Δhinge is predicted to be completely coiled-coil and therefore presumably rigid (compare to full-length stalk in Figure 1A). c) KLP-18 stalk flexibility tested in size exclusion chromatography experiment. MBP-stalk applied to a size exclusion column in high salt (300mM, blue) and low salt (20mM, gold) buffer. Indicated fractions probed for KLP-18 in a Western blot then quantified. Mean +/- sd of percent of total band intensity is shown. High salt: n = 3 over 2 purifications, low salt: n = 2 over 1 purification.

**Figure S3:**
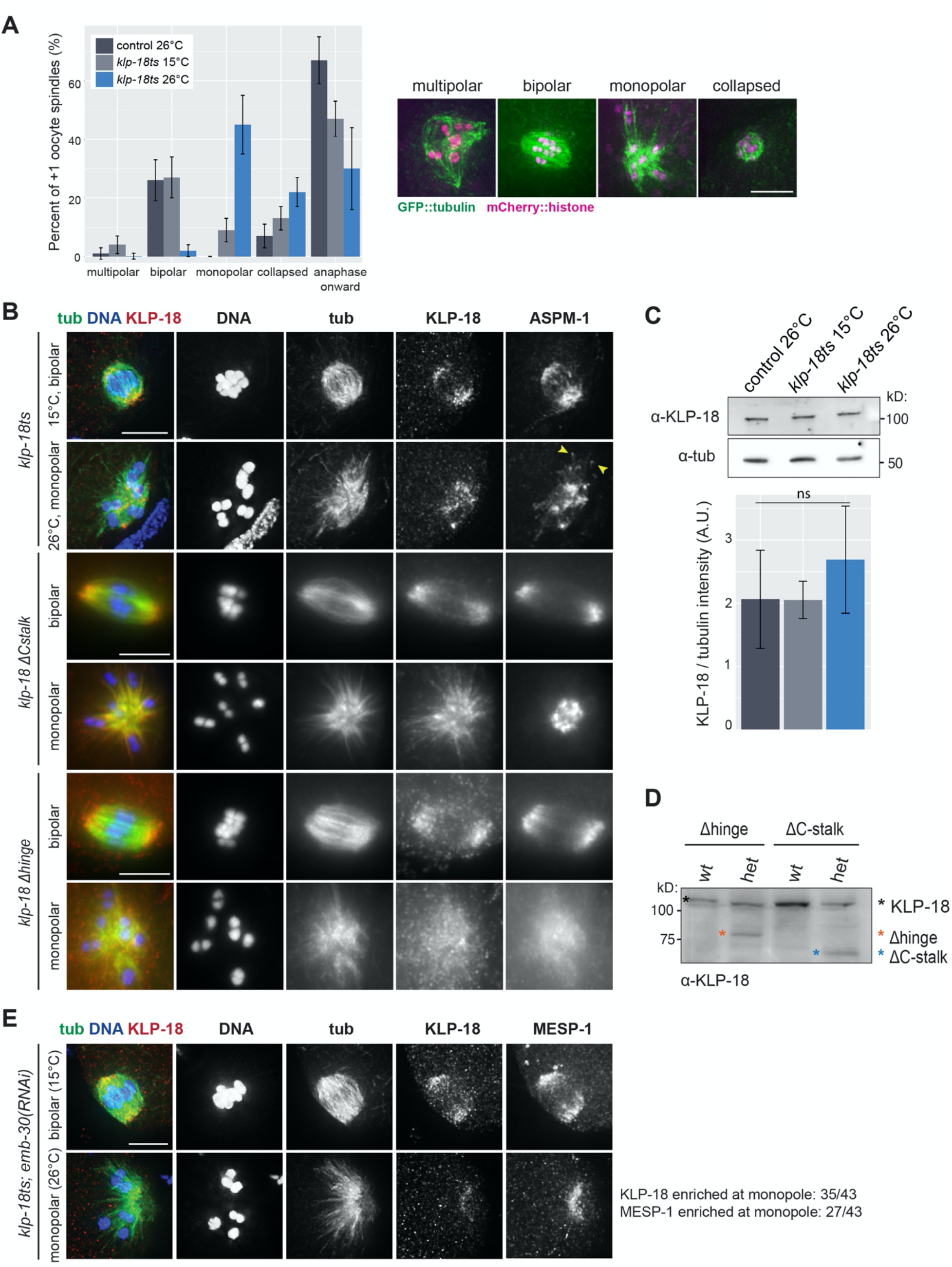
KLP-18 mutant strains express truncated protein and result in monopolar spindles. a) Oocyte spindle morphology in the most recently fertilized (+1) embryo of control (dark gray) and *klp-18ts* worms expressing GFP::tubulin and mCherry::histone. During normal spindle assembly, minus ends are sorted away from chromosomes, to form multiple poles (“multipolar”) that coalesce to form a bipolar spindle (“bipolar”) [12]. If minus ends are not sorted outwards, a monopolar spindle forms with chromosomes arranged in a rosette (“monopolar”) [9–12]; these chromosomes then move inwards in anaphase (“collapsed”) [51]. Spindle morphology in *klp-18ts* was quantified at permissive (15°C, light gray) and restrictive (26°C, blue) temperatures; spindles that had entered anaphase (with two sets of separated chromosomes) and Meiosis II spindles were grouped together as “anaphase onwards”. Note that this data is from the same quantification depicted in Figure 2B, but in this case all categories are represented, including spindles that had progressed to anaphase (in Figure 2B the “anaphase onwards” category and the “collapsed” category, representing monopolar anaphase, were excluded to emphasize the spindle assembly phenotype). Bars represent mean percentage +/- sd. For *klp-18ts* conditions, n = 4 experiments; for control, n = 3 experiments. b) DNA (blue), tubulin (green), KLP-18 (red), and ASPM-1 (not in merge) localization in *klp-18(or447ts), klp-18ΔC-stalk*, and *klp-18Δhinge* mutant worms. Representative images of wild type bipolar and mutant monopolar spindles shown for each strain. In the *klp-18ts* mutant, there are some ASPM-1 foci on the outside of the monopolar spindle (arrowheads), suggesting that there is weak microtubule sorting activity in this mutant that allows some minus ends to be pushed outwards. Images for *klp-18ts* are deconvolved to better show ASPM-1 foci on the outside of the aster; other images are not deconvolved. Scale bars = 5μm. c) Western blot of control (dark gray) and *klp-18ts* worms at permissive (15°C, light gray) and restrictive (26°C, blue) temperature probed with an anti-KLP-18 antibody. Representative blot (top) with quantification of normalized KLP-18 band intensity (below, mean +/- sd). Quantified KLP-18/tubulin intensity was not significantly different between any of the conditions over n = 3 experiments (p > 0.05, paired one-tail Student’s t-Test). d) Western blot of progeny from homozygous wild type (‘wt’) and heterozygous mutant (‘het’) *klp-18Δhinge* or *klp-18ΔC-stalk* parents probed with an anti-KLP-18 antibody. Wild type (black), *Δhinge* (orange), and *ΔC-stalk* (blue) bands indicated by asterisks. e) DNA (blue), tubulin (green), KLP-18 (red), and MESP-1 (not in merge) localization in *klp-18ts* worms at 15°C and 26°C; *emb-30(RNAi)* was used to induce metaphase I arrest. These images are deconvolved. Quantification shown to the right of the images; for details on how enrichment was defined see “Figure Quantification” section of Materials and Methods. Scale bar = 5μm.

**Figure S4:**
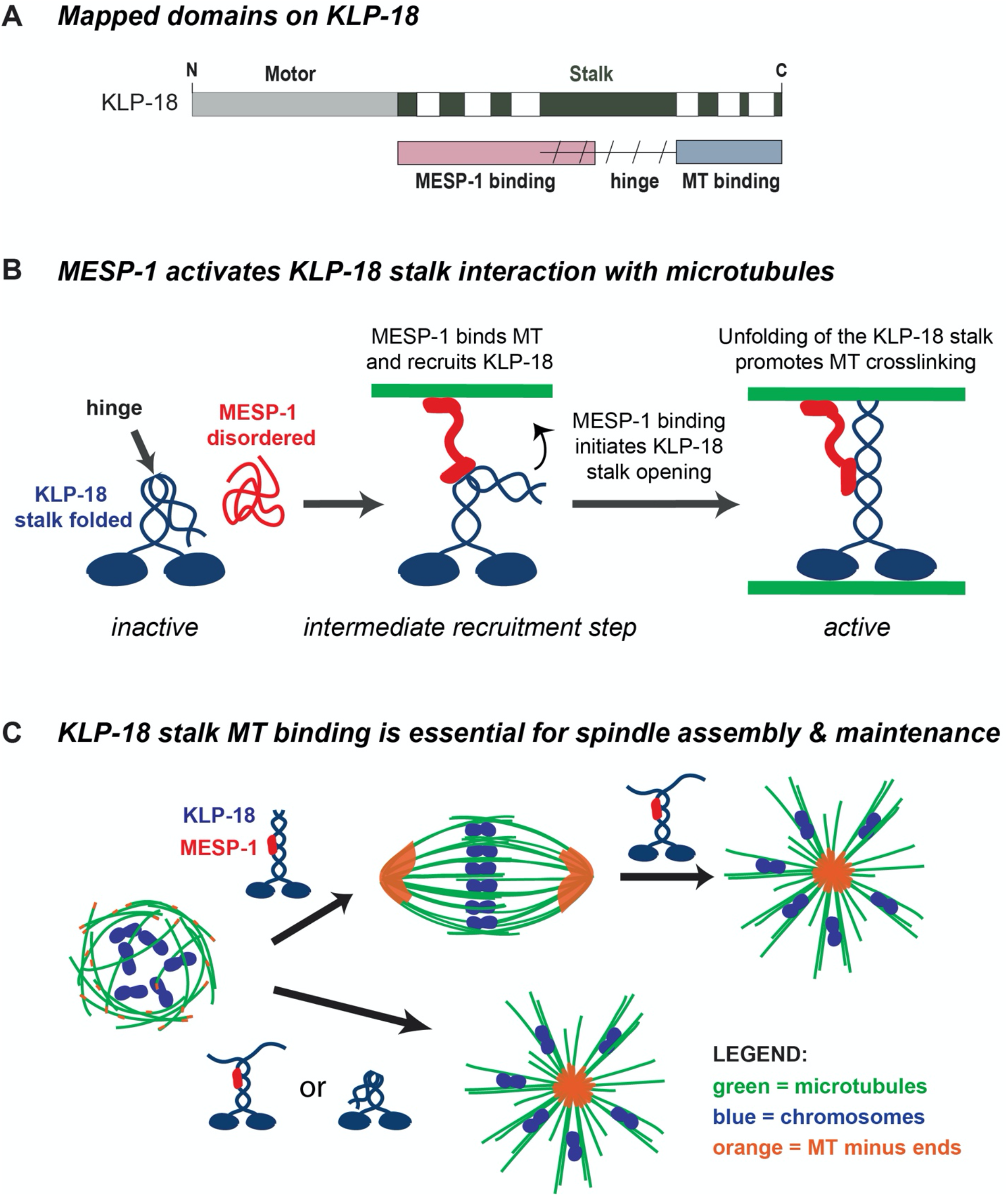
Model. a) KLP-18 stalk contains a MESP-1 binding site in its N-terminal half (pink), a microtubule binding site in its C-terminal half (blue), and likely contains a flexible hinge region (slanted dashed lines) that functions to self-inhibit the motor. The MESP-1 binding site and the hinge region overlap to some extent, as shown. b) Before complex formation, KLP-18 is in an auto-inhibited inactive state and MESP-1 is disordered and non-functional. Upon complex formation, MESP-1 targets KLP-18 to spindle microtubules and promotes KLP-18 stalk opening. Once KLP-18 is activated through unfolding, the C-terminal microtubule binding site can bind to microtubules, allowing the KLP-18/MESP-1 complex to crosslink microtubules and promote spindle bipolarity. After the initial targeting of the complex, MESP-1 may remain bound to one of the two crosslinked microtubules. In this simplified model, we show MESP-1 bound to the microtubule associating with the KLP-18 C-terminal binding site, but it is also possible that MESP-1 could bind the microtubule associated with the KLP-18 motor domain. c) Disruption of the KLP-18 C-terminal microtubule binding site impairs microtubule sorting during spindle assembly and leads to the collapse of pre-formed bipolar spindles. Both the C-terminal microtubule binding site and interaction with MESP-1 are essential for KLP-18 function; MESP-1 is likely essential for initial targeting to spindle microtubules and the C-terminal microtubule binding site is essential for motor function. KLP-18 is shown as blue and MESP-1 is shown as red in the cartoon. On the spindle diagrams, microtubules are green, chromosomes are blue, and microtubule minus ends are orange.

**Movie S1. Metaphase I-arrested oocyte spindle shortens and then forms a monopole upon temperature shift in *klp-18(or447ts)* mutant**.

Representative movie of spindle reorganization described and quantified in Figures 4B and 4C. Chromosomes (mCherry::histone, magenta) and microtubules (GFP::tubulin, green) visualized in an *emb-30(RNAi) klp-18(or447ts)* oocyte dissected then shifted to the restrictive temperature. Scale bar = 10μm.

